# Artificial selection finds new hypotheses for the mechanism of *Wolbachia*-mediated dengue blocking in mosquitoes

**DOI:** 10.1101/847087

**Authors:** S.A. Ford, I. Albert, S. L. Allen, S. Chenoweth, M. Jones, C. Koh, A. Sebastian, L.T. Sigle, E.A. McGraw

**Affiliations:** Huck Institutes for the Life Sciences, Penn State University, University Park, 16802, PA, USA; School of Biological Sciences, Monash University, Melbourne, Vic, 3800, Australia; Department of Zoology, University of Oxford, OX1 3SZ, England, UK; School of Biological Sciences, The University of Queensland, St. Lucia, Queensland, Australia; Institut für Populationsgenetik, Vetmeduni Vienna, Vienna, Austria; Department of Virology, Institut Pasteur, Paris, 75015, France

**Keywords:** Disease control, *Aedes aegypti*, *Wolbachia pipientis*, genetic variation, evolution

## Abstract

*Wolbachia* is an intracellular bacterium that blocks virus replication in insects and has been introduced into the mosquito, *Aedes aegypti* for the biocontrol of arboviruses including dengue, Zika and chikungunya. Despite ongoing research, the mechanism of *Wolbachia*-mediated virus blocking remains unclear. We recently used experimental evolution to reveal that *Wolbachia*-mediated dengue blocking could be selected upon in the *A. aegypti* host and showed evidence that strong levels of blocking could be maintained by natural selection. In this study, we investigate the genetic variation associated with blocking and use these analyses to generate testable hypotheses surrounding the mechanism of *Wolbachia*-mediated dengue blocking. From our results, we hypothesise that *Wolbachia* may block virus replication by increasing the regeneration rate of mosquito cells via the Notch signalling pathway. We also propose that *Wolbachia* modulates the host’s transcriptional pausing pathway either to prime the host’s anti-viral response or to directly inhibit viral replication.

## Introduction

The *Aedes aegypti* mosquito transmits arboviruses that cause morbidity and mortality globally, including dengue, Zika and chikungunya (Kraemer et al. 2019). *Wolbachia* are vertically-transmitted, intracellular alpha-proteobacteria that can protect insects from viruses, termed virus ‘blocking’ (Teixeira, Ferreira, and Ashburner 2008, Hedges et al. 2008). *Wolbachia* have been artificially introduced into *A. aegypti*, where they are stably-inherited (Gloria-Soria, Chiodo, and Powell 2018) and block the transmission of dengue, Zika and chikungunya viruses (Dutra et al. 2016, Tan et al. 2017, Ye et al. 2015, Moreira et al. 2009, Aliota, Peinado, et al. 2016, Aliota, Walker, et al. 2016). *Wolbachia* have since been spread through natural populations of mosquitoes where they have reduced the incidence of locally-transmitted dengue (Hoffmann, Ross, and Rašić 2015, Hancock et al. 2016, Walker et al. 2011, O’Neill 2018, Jiggins 2017, Schmidt et al. 2017, O’Neill et al. 2018).

Despite substantial discoveries, the mechanism of *Wolbachia*-mediated virus blocking remains unclear. Research using insect cells suggests that blocking impacts early viral genome replication (Rainey et al. 2016, Thomas et al. 2018). Flaviviruses use the host cytoskeleton and alter the endoplasmic reticulum (ER) to form sites of replication, termed replication complexes (RCs) (Lindsey et al. 2018). *Wolbachia* also modify the ER and rely upon it for protein degradation (Geoghegan et al. 2017, Molloy et al. 2016, White et al. 2017). Moreover, *Wolbachia* modulates the cytoskeleton to ensure their localization in the cell during mitosis and meiosis (Lindsey et al. 2018). Together, flaviviruses and *Wolbachia* are likely to compete for the ER and cytoskeleton, along with resources (Caragata et al. 2013). Other pathways have also been shown to affect *Wolbachia*-mediated blocking but do not seem to be required, e.g. host immune priming (Rancès et al. 2012, Rancès et al. 2013), the RNAi pathway (Terradas, Joubert, and McGraw 2017), and host miRNAs (Zhang et al. 2013, Hussain et al. 2011, Rainey et al. 2016), the production of reactive oxygen species (ROS) (Pan et al. 2012, Pan et al. 2018, Wong, Brownlie, and Johnson 2016) and XRN1-mediated virus degradation (Thomas et al. 2018).

Recently, we artificially selected upon *Wolbachia*-infected *A. aegypti* mosquitoes (*w*Mel.F strain) to test if we could increase or decrease the strength of *Wolbachia*-mediated dengue blocking (Ford et al. 2019). We were able to isolate mosquitoes that had lost 45% of *Wolbachia*-mediated blocking compared to randomly-selected control populations and identified that this resulted from genetic variation in *A. aegypti* and not *Wolbachia*. Interestingly, we could not increase blocking strength and found evidence to suggest that genotypes exhibiting strong levels of blocking were already at a high frequency due to an inherent fitness advantage. These data indicate the potential for natural selection to maintain blocking. In addition to this, these data provide a great opportunity to gain insight into the mechanism of *Wolbachia*-mediated dengue blocking. In this previous study, we investigated two candidate genes that had undergone large changes in allele frequencies as a result of selection on blocking. We found one gene whose expression increased with blocking strength. This gene encoded a cadherin protein (AAEL023845). Cadherins are cell-cell adhesion proteins that mediate cell signalling and intracellular trafficking (Yamagata, Duan, and Sanes 2018), yet it remains unclear how this gene may be involved in *Wolbachia*-mediated dengue blocking.

In this study, we investigate 61 *A. aegypti* genes that exhibited significant changes in allele frequencies between the mosquito populations that we selected for high and low *Wolbachia*-mediated dengue blocking (Ford et al. 2019). We then used these data to generate novel and testable hypotheses surrounding the mechanism of *Wolbachia*-mediated dengue blocking. We found that genes under selection for blocking were significantly enriched for signal transduction and transcription regulation. More specifically, we found that genes involved in neurogenesis, the Notch signalling pathway and cell-cell adhesion are the most commonly selected upon, leading to the hypothesis that Notch-mediated cell replenishment may be important for *Wolbachia*-mediated viral blocking. We also revealed that the host’s transcriptional pausing pathway could be involved in blocking and find evidence for the host’s oxidative stress response and cytoskeleton, consistent with previous studies.

## Results and Discussion

### Gene ontology (GO) term enrichment analysis

We investigated 61 *A. aegypti* genes that showed significant changes in allele frequencies between the mosquito populations selected for high and low *Wolbachia*-mediated blocking (Supplementary File 1) in our previous study (Ford et al. 2019). We explain the generation of these significant genes in the Materials and Methods. We tested for enriched gene ontology (GO) terms using the Singular Enrichment Analysis (SEA) available on AgriGo v2 for the AaegL3.3 locus ID (VectorBase) (Fig. 1 and Supplementary File 2, generated by AgriGo v2). The most specific GO terms (as measured by “Bgitem”, which is the total number of genes annotated with that GO term in the *A. aegypti* genome) that were significantly enriched included: ‘regulation of gene expression’ (Fisher test with Benjamini-Hochberg false discovery rate, FDR, P-value adjustments P=0.018), ‘nucleobase-containing compound metabolic process’ (FDR-corrected P=0.0113), ‘cellular macromolecule metabolic process’ (FDR-corrected P=0.0186) and ‘signal transduction’ (FDR-corrected P=0.0174).

**Figure 1.**
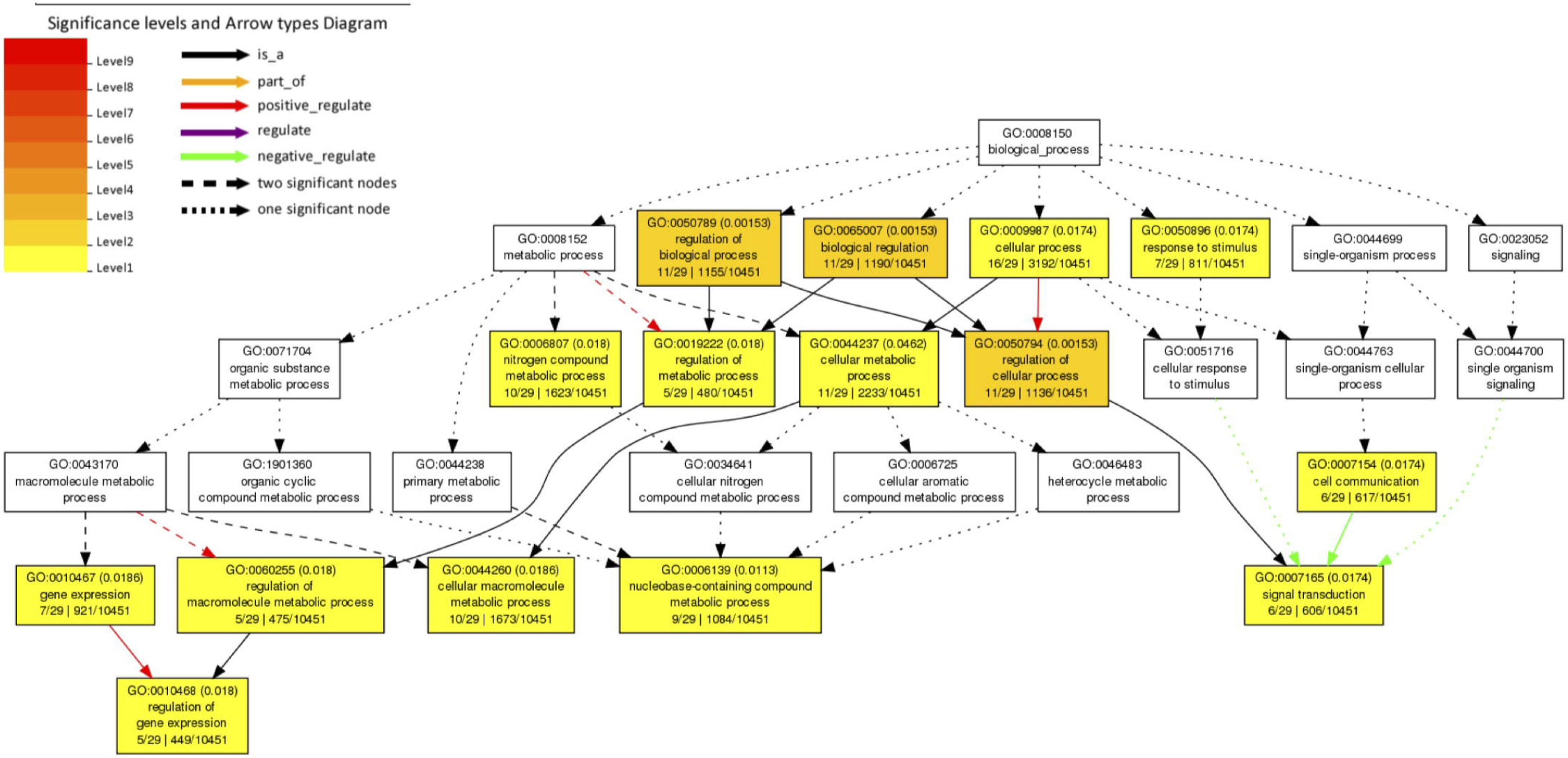
Gene Ontology (GO) term enrichment analysis of *A. aegypti* genes containing SNPs associated with *Wolbachia*-mediated dengue blocking. We performed a Singular Enrichment Analysis (SEA) using AgriGo v2 on 61 *A. aegypti* genes that contained SNPs that were significantly differentiated between mosquito populations selected for high vs. low *Wolbachia*-mediated dengue blocking (Supplementary File 1) in our previous study (Ford et al. 2019). The SEA utilised a Fisher test with Benjamini-Hochberg (FDR) P-value adjustments for multiple comparisons. P-values are shown in brackets. GO terms and the gene entries assigned to them are listed in Supplementary File 2.

### Cell replenishment

We then grouped the genes by function, using the OrthoDB, Uniprot and Interpro databases (Table 1). Consistent with the significant GO term ‘signal transduction’, we found genes for neurogenesis, the Notch signalling pathway, cell cycle and cell-cell adhesion, including the previously described cadherin gene (AAEL023845). Together, this gene profile suggests that cell replenishment pathways could be important for *Wolbachia*-mediated dengue blocking. It has been found that the Notch signalling pathway confers resistance to the dengue virus in *A. aegypti* by controlling host cell regeneration rate whereby mosquitoes with faster cell regeneration in the midgut are significantly more resistant to the dengue virus (Taracena et al. 2018). Moreover, the depletion of cadherin proteins is known to downregulate the Notch signalling pathway and so cell replenishment (Hatakeyama et al. 2014, Sasaki et al. 2007).

**Table 1.** Functional groups of annotated genes under selection for *Wolbachia*-mediated dengue blocking. *A. aegypti* genes that contain single nucleotide polymorphisms (SNPs) that were significantly differentiated between mosquito populations selected for high vs. low *Wolbachia*-mediated dengue blocking (Ford et al. 2019). The genes are listed if they have been annotated and are grouped based upon their function according to OrthoDB, Uniprot and Interpro databases. The genes are listed by the number of significantly differentiated SNPs (see Materials and Methods). P-values are listed in Supplementary File 1.

| No. of SNPs | Gene ID | Gene name (*D. melanogaster* orthologue) | Putative function |
| --- | --- | --- | --- |
| **Neuronal function** |  |  |  |
| 17 | AAEL004396 | Dopamine receptor D3 | Synaptic transmission |
| 12 | AAEL004389 | Alpha-mannosidase | N-linked glycosylation pathway in Golgi. Linked to embryonic development and the positive regulation of neurogenesis |
| 6 | AAEL002769 | Homeo-prospero domain | Neural differentiation. Transcription factor protein that is expressed in all neural lineages of *Drosophila* embryos |
| 5 | AAEL009466 | Protein Kinase C | Regulation of neurotransmitter release, ion channels, growth and differentiation, and neural plasticity |
| 1 | AAEL002876 | CUB domain | Inflammation, neurotransmission, receptor-mediated endocytosis, axon guidance and angiogenesis, cell signalling |
| 1 | AAEL006120 | G protein-coupled receptor kinase | Regulation of receptor signalling, particularly in neurons. |
| **Notch signalling** |  |  |  |
| 6 | AAEL008617 | Hairy | bHLH transcriptional repressor involved in embryonic segmentation and peripheral neurogenesis |
| 3 | AAEL014528 | SNW domain-containing protein 1 | Positive regulation of neurogenesis; positive regulation of transcription of Notch receptor target |
| 3 | AAEL010513 | Helt bHLH transcription factor | Central nervous system development, Notch signalling pathway |
| 1 | AAEL006817 | Achaete-scute homolog 1 | Neuronal differentiation. Notch signalling pathway. bHLH transcription factor binding |
| **Cell-cell adhesion** |  |  |  |
| 90 | AAEL023845 | Cadherin(cad87A) | Cadherin and neurexin bind trans-synaptically |
| 19 | AAEL008334 | Afadin (canoe) | Organisation of cell junctions during embryogenesis |
| 5 | AAEL004233 | igLON family member 5 | Cell adhesion molecule |
| 4 | AAEL021067 | Immunoglobulin-like domain | Cell-cell recognition |
| 2 | AAEL019752 | Neurexin 3 isoform X1 | Cadherin and neurexin bind trans-synaptically |
| 2 | AAEL010721 | Leucine-rich repeat-containing protein 4C | May promote neurite outgrowth of developing thalamic neurons. Synaptic membrane adhesion. Regulation of axonogenesis |
| 2 | AAEL006191 | Rho GTPase-activating protein 39 isoform X1 | Postsynapse organisation |
| 1 | AAEL010881 | Netrin receptor UNC5D | Cell-cell adhesion via plasma-membrane adhesion molecules |
| 1 | AAEL007205 | Gata-binding factor-c (Grain) | Grain is a transcription factor from the GATA family. It regulates the expression of receptors and adhesion molecules such as unc-5 and Fas2 involved in axon guidance during development |
| **Ion channels** |  |  |  |
| 1 | AAEL025294 | Otopetrin-1 | Proton channel activity |
| 1 | AAEL018306 | Calcium activated potassium channel subunit alpha-1 | Membrane depolarization |
| 1 | AAEL021538 | Chloride channel accessory 1 | Transmembrane transfer of chloride |
| **Cell cycle** |  |  |  |
| 2 | AAEL011522 | Ras-related and estrogen-regulated growth inhibitor | Negative regulation of cell growth |
| 2 | AAEL014044 | Protein Skeletor, isoforms B/C | Spindle assembly, cell division |
| 1 | AAEL005802 | Structural maintenance of chromosomes protein | Essential for successful chromosome transmission during replication and segregation of the genome in all organisms |
| 1 | AAEL004700 | Cyclin-dependent kinase-like 1 | Cell division control |
| 1 | AAEL004572 | Ligand-dependent nuclear receptor corepressor-like protein (Ecdysone-induced protein 93F) | Transcription factor regulating cell death |
| **Transcriptional pausing** |  |  |  |
| 12 | AAEL018695 | metastasis-associated protein MTA | Heterochromatin organization involved in chromatin silencing |
| 3 | AAEL005813 | Negative elongation factor A | Negative regulation of transcription by RNA polymerase II. An essential role in postembryonic development. Transcriptional pausing controls rapid antiviral innate immune response |
| 2 | AAEL024508 | Transcription elongation factor B polypeptide 3 (Elongin A) | Positive regulation of transcription elongation. Suppresses the transient pausing of RNA polymerase II |
| 2 | AAEL019422 | Histone deacetylase | Alters transcription |
| 1 | AAEL024283 | BTB/POZ domain | Transcriptional regulators that are thought to act through the control of chromatin structure |
| **Cytoskeleton/transport** |  |  |  |
| 5 | AAEL018215 | Dedicator of cytokinesis protein 2 | Actin cytoskeleton organization |
| 4 | AAEL002173 | TRAF3-interacting protein 1 | Regulation of microtubule cytoskeleton organization. Negative regulation of defense response to virus |
| 2 | AAEL002571 | Band 4.1-like protein | Cytoskeletal protein binding. actomyosin structure organization |
| 1 | AAEL019696 | Unconventional myosin-VIIb | Actin-based motor molecules with ATPase activity. Cell differentiation |
| 1 | AAEL014511 | Zinc finger, AN1-type | Positive regulation of intracellular protein transport |
| **Oxidative stress** |  |  |  |
| 4 | AAEL013026 | Guanylate cyclase 1 soluble subunit alpha 1 | Nitric oxide mediated signal transduction. Response to oxidative stress |
| 1 | AAEL019639 | Haem peroxidase | Reduction of hydrogen peroxide. Response to oxidative stress |
| **Endoplasmic Reticulum (ER) functioning** |  |  |  |
| 1 | AAEL024147 | ELO family | Small molecule metabolism. Likely located on the endoplasmic reticulum. |
| 1 | AAEL006208 | DDB1- and CUL4-associated factor 15 | Protein ubiquitination. *Wolbachia* relies on it for protein degradation |
| **RNA editing** |  |  |  |
| 3 | AAEL013723 | Heterogeneous nuclear ribonucleoprotein L-like | Regulator of alternative splicing |
| 3 | AAEL027527 | tRNA-specific adenosine deaminase 1 (ADAR) | tRNA processing. It is involved in behaviour, glucose metabolism and response to several stresses such as heat, hypoxia and oxidative stress. Prevents the proper translation of viral RNAs |
| 1 | AAEL019447 | Intron-binding protein aquarius | Splicing factor |
| 1 | AAEL020230 | Zinc finger CCCH domain-containing protein 13 isoform X1 | Methylation and splicing of RNAs. Controls embryonic stem cells (ESCs) pluripotency |
| 1 | AAEL006197 | Zinc finger CCHC-type and RNA-binding motif-containing protein 1 | RNA splicing |

We therefore hypothesise that *Wolbachia* could block virus by increasing cell-adhesion and host cell regeneration rate via Notch signalling. This hypothesis would be consistent with our previous observation that the reduction in viral blocking was associated with reduced expression of the cell-cell adhesion gene, cadherin (Ford et al. 2019). Slower cell regeneration would also explain the inherent fitness cost that we observed in mosquitoes with weaker viral blocking (Ford et al. 2019). How *Wolbachia* might alter this pathway is unknown, but *Wolbachia* has been found to stimulate mitosis in nematodes (Foray et al. 2018) and increase mosquito metabolism (Evans et al. 2009).

It is unclear whether neurogenesis specifically plays a role in *Wolbachia*-mediated blocking or whether the gene annotations exhibit a bias from *Drosophila* developmental studies and these genes instead function in cell proliferation and differentiation more generally (Taracena et al. 2018). The Zika virus has, however, been found to dysregulate neurogenesis in humans through the Notch signalling pathway (Ferraris et al. 2019) and by degrading adherens junction proteins (Yoon et al. 2017). Although this has not been shown with the dengue virus, it does upregulate the expression of cell-cell junction genes in humans (Afroz et al. 2016). *Wolbachia* is also known to cause neuropathology (Min and Benzer 1997, Moreira et al. 2011) and control host factors during neurogenesis (Albertson et al. 2009) 2009). Critically, if *Wolbachia*-mediated virus blocking involves pathways by which viruses cause neuropathology in humans then it could select for viruses with altered disease severity. This exemplifies the importance of understanding the mechanism of blocking.

### Transcriptional pausing

Consistent with the significant GO term ‘regulation of gene expression’, we found a number of genes involved in transcriptional pausing. The transcriptional pausing pathway plays a role in the *Drosophila* antiviral response by priming the rapid transcription of immune genes involved in RNA silencing, autophagy, JAK/STAT, Toll and Imd pathways along with components of Toll receptors (Xu et al. 2012). Transcriptional pausing can also directly restrict viral transcription in mammalian cells (Fujinaga et al. 2004, Rao et al. 2006). Transcriptional pausing involves Negative and Positive Elongation Factors (NELF and P-TEFb) and is associated with open chromatin structures. Here, we find genetic variation associated with blocking in genes encoding NELF and P-TEFb (AAEL005813 and AAEL024508, respectively) and genes involved in chromatin-structure mediated silencing (AAEL018695, AAEL024283 and AAEL019422). We hypothesise that *Wolbachia*-mediated viral blocking could trigger the host’s transcriptional pausing pathway to prime the antiviral response or to directly inhibit virus transcription.

### Cytoskeleton/transport

The dengue virus utilises the cytoskeleton to enter/exit the cell and recruit resources to RCs (Lindsey et al. 2018). Simultaneously, *Wolbachia* utilise the cytoskeleton for successful transmission during mitosis and meiosis and so may interfere with the virus life-cycle (Lindsey et al. 2018). In our dataset, we identified many genes with putative links to the cytoskeleton, including genes related to microtubules (AAEL002173), actin (AAEL018215) and associated motors (AAEL019696). Of particular interest is the gene TRAF3-interacting protein 1 (AAEL002173). TRAF3 has been implicated in an antiviral response in the sand fly, *Lutzomyia longipalpis* (Martins-da-Silva et al. 2018).

### Oxidative stress

Oxidative stress is a by-product of *Wolbachia* infection and can trigger anti-microbial host responses (Lindsey et al. 2018). We find variation associated with blocking in a gene that reduces hydrogen peroxide (AAEL019639) and in another that responds to nitric oxide (AAEL013026). Changes in these genes could either: remove ROS, e.g. to enable the host to tolerate *Wolbachia* infection; or prevent the removal of ROS, e.g. to trigger an anti-microbial response, or stimulate mitosis and so increase cell regeneration (Taracena et al. 2018).

## Conclusion

From reviewing *A. aegypti* genes isolated by artificial selection, we have been able to present new hypotheses for the mechanism of *Wolbachia*-mediated dengue blocking. Signal transduction and the regulation of transcription were significantly enriched in this dataset, corresponding to large sets of genes for cell regeneration and the transcriptional pausing pathway. Our data led to two main hypotheses: 1) efficient regeneration of mosquito cells increases resistance against flaviviruses and *Wolbachia* could promote this by modulating adherens junctions and Notch signalling; and 2) *Wolbachia* could trigger the host’s transcriptional pausing pathway to either prime the hosts’ anti-viral response or inhibit viral transcription. Consistent with previous studies, we also found evidence that oxidative stress and the cytoskeleton are likely to be involved in blocking.

## Materials and Methods

### Ethics

The evolution experiment was carried out at Monash University in Melbourne. The Monash University Human Research Ethics Committee gave ethical approval for human volunteers to provide blood meals to mosquitoes not infected with dengue virus (permit: CF11/0766-2011000387). One volunteer was involved throughout the study and provided written consent.

### Mosquitoes

For the evolution experiment we used a population of *A. aegypti* that were infected with the wMel.F strain of *Wolbachia* (Walker et al. 2011, Hoffmann et al. 2011)and had been maintained in the laboratory for 33 generations. Every three generations, they were outcrossed with *Wolbachia*-free mosquitoes collected from Queensland, Australia, to maintain standing genetic variation (Terradas et al. 2017, Hoffmann et al. 2011). Mosquitoes collected for outcrossing were replaced every six generations. During outcrossing, ~30% of males from the laboratory population were replaced with males from the collected population. The wild-type populations used in this study to measure gene expression were the AFM line obtained from Zhiyong Xi, collected 12 months prior by Pablo Manrique in Merida, Mexico.

### Dengue virus

For the evolution experiment, we used dengue virus serotype 3, isolated from Cairns(Ye et al. 2016, Ritchie et al. 2013). Virus was grown within C6/36 Aedes albopictus cells following standard methods(Terradas et al. 2017). Cells were grown to 80% confluency at 26 °C in T175 tissue culture flasks containing 25 ml RPMI 1640 media (Life Technologies) supplemented with 10% foetal bovine serum (Life Technologies), 2% HEPES (Sigma–Aldrich) and 1% Glutamax (Life Technologies). The media was then replaced with 25 ml RPMI supplemented with 2% foetal bovine serum, 2% HEPES and 1% Glutamax, and 20 μl virus was added. After 7 days, cells were scraped off and the suspension was centrifuged at 3,200g for 15 min at 4 °C. The supernatant was frozen in single-use aliquots at −80 °C, and all experiments were conducted using these aliquots. Virus titres were measured from a thawed aliquot by: (1) mixing 20 μl with 200 μl of TRIzol (Invitrogen); (2) extracting the RNA following the manufacturer’s protocol and treating with DNAse I (Sigma–Aldrich); and (3) quantifying dengue virus RNA using quantitative reverse-transcription PCR (RT-qPCR) (see ‘Dengue virus quantification’). Three independent extractions were performed and two replicates of each extraction were measured to generate an average value of 1.80 × 106 genomic copies of the dengue virus per ml.

### Dengue virus quantification

Dengue virus was quantified via RT-qPCR using the LightCycler 480 (Roche). We used the TaqMan Fast Virus 1-Step Master Mix (Thermo Fisher Scientific) in a total reaction volume of 10 μl, following the manufacturer’s instructions(Terradas et al. 2017). The list of primers and probes is given in Supplementary Table 1. The temperature profile used was: 50 °C for 10 min; 95 °C for 20 s; 35 cycles of 95 °C for 3 s; 60 °C for 30 s; 72 °C for 1 s; and 40 °C for 10 s. Data were analysed using absolute quantification where the dengue virus copy number per sample was calculated from a reference curve. This reference curve was made up from known quantities of the genomic region of the virus that the primers amplify. This genomic region had previously been cloned into the pGEM-T plasmid (Promega) and transformed into Escherichia coli(Ye et al. 2014). After growing E. coli in liquid Luria broth (LB) overnight at 37 °C, we extracted the plasmid using the PureYield Plasmid Midiprep System kit (Promega) and linearized it by restriction digest. We then purified the plasmid using phenol-chloroform extraction, resuspended in 20 μl of UltraPure distilled water (Invitrogen) and quantified it by Qubit. A dilution series of 107, 106, 105, 104, 103, 102 and 101 copies of the genomic fragment was created and frozen as single-use aliquots. All assays measuring viral load used these aliquots, and three replicates of the dilution series were run on every 96-well plate to create a reference curve for dengue virus quantification.

### Evolution experiment

We selected for low and high *Wolbachia*-mediated dengue blocking alongside a control treatment where mosquitoes were selected at random. Each treatment included three independent populations generated from an ancestral population of mosquitoes using a random number generator(Kawecki et al. 2012). For each generation, eggs were hatched in trays (30 cm × 40 cm × 8 cm) containing 21 of autoclaved reverse osmosis water to achieve 150–200 larvae per tray. Larvae were fed ground TetraMin tablets and reared under controlled conditions of temperature (26 ±2 °C), relative humidity (~70%) and photoperiod (12 h:12 h light:dark). After pupation, pupae were placed within 30 cm × 30 cm × 30 cm cages in cups containing autoclaved reverse osmosis water for eclosion to achieve ~450 mosquitoes per cage. Mosquitoes were fed 10% sucrose water from dental wicks. When mosquitoes were 5–7 d old, each population was allowed to blood-feed on a human volunteer in a random order. Females that fed were separated into cups enclosed with mesh that contained moist filter paper to provide an oviposition site. Mosquitoes were fed 10% sucrose water from cotton wool.

After 4 d, eggs were collected, numbered and dried following a standard protocol for short-term egg storage(Zheng et al. 2015). The number of each set of eggs was written on the cups of the corresponding female. Egg collection was done before infection with dengue to prevent vertical transmission of the virus (Joshi, Mourya, and Sharma 2002). Between 40 and 70 females from each of the high- and low-blocking populations were anaesthetized with CO_2_, injected with 69 nl of the dengue virus stock (equalling ~124 genomic copies of dengue; see ‘Dengue virus’) and returned to their numbered cups. Virus was delivered at a speed of 46 nl s−1 into the thorax using a pulled glass capillary needle and a manual microinjector (Nanoject II; Drummond Scientific). This controlled the infection dose by removing the variation that would have resulted from oral feeding, to ensure successful artificial selection. This method also ensured a sufficient number of infected mosquitoes to select between.

At 7 d post-infection, females were anaesthetized with CO_2_, placed into individual wells of 96-well plates containing 50 μl extraction buffer and homogenized with a 3-mm glass bead. The extraction buffer was made up of squash buffer (10 mM Tris (pH 8.2), 1 mM EDTA and 50 mM NaCl) (Yeap et al. 2014) with proteinase K at a concentration of 12.5 μl ml−1 (Bioline). Samples were then incubated for 5 min at 56 °C and 5 min at 95 °C. We then measured the viral load per mosquito using RT-qPCR (see ‘Dengue virus quantification’). This method was used for rapid phenotype determination of a large number of samples. Mosquitoes were then ranked in order: (1) from the lowest viral load in the high-blocking populations; (2) from the highest viral load in the low-blocking populations; and (3) using a random number generator in the random population. Eggs from each mosquito were hatched into separate cups of autoclaved reverse osmosis water. The next day, larvae were taken from cups in rank order until ~200 larvae were collected for each replicate population. On average, offspring were taken from six mosquitoes per replicate population per generation. This was done to impose the strongest selection pressure possible while ensuring that enough mosquitoes would be reared for selection in the subsequent generation. At this point, the protocol was repeated. In total, four rounds of selection were completed.

### Genomic analysis

DNA was extracted from 90 individual mosquitoes from the ancestral population and each evolved population after 4 generations of selection. We extracted DNA from the TRIzol reagent (Invitrogen) using a modified version of the manufacturer’s protocol with additional washing steps using phenol, chloroform and isoamylalcohol. DNA was sequenced using an Illumina HiSeq 3000 with 150-base pair paired-end reads.

FastQC version 0.11.4 was used with default settings to check the quality of the raw reads. To minimize false positives, Trimmomatic version 0.36 was used to trim the 3’ ends if the quality was <20, and the reads were discarded if trimming resulted in reads that were <50 base pairs in length (<0.5%). We mapped the resulting reads to the *A. aegypti* assembly (Liverpool AGWG-AaegL5) using BWA MEM 2.2.1, and checked for quality using Qualimap version 2.2.1. Indel realignment was completed using GATK version 3.8.0. Duplicates were removed using Picard version 2.17.8, and reads with poor mapping quality were removed using SAMtools 1.6 and filtering via hex flags: -q 20 (only include reads with a mapping quality of ≥ 20); -f 0 × 002 (only include reads with all of the flags mapped in a proper pair); -F 0 × 004 (only include reads with none of the flags unmapped); and -F 0 × 008 (only include reads with none of the flags mate unmapped). Around 10% of reads were PCR duplicates (615,305,021) and ~58% of reads failed mapping quality filters (3,655,353,869). The quality was checked using Qualimap.

SNPs were called using PoPoolation2 based on a minimum coverage of 20 and a maximum coverage of 200. Coverage was ~46 after duplicate and low-quality mapping were removed (~51 before). We ran an alternative method to call variants to cross-check the output from the above method. Variants were called for each sample using the GATK HaplotypeCaller tool (gatk-4.0.8.1) using default settings except for ploidy, which was set to 10. A multi-sample variant file was then created merging the vcf files using the ‘bcftools merge’ command. SNPs from the original analysis were retained if at least one population had the same SNP called by GATK.

We identified SNPs that were significantly differentiated between treatments (see ‘Statistics’) and annotated them with gene information using gene transfer format files and bedtools intersect (bedtools version 2.25). Annotation files were downloaded from VectorBase (AaegL5.1). Information on *A. aegypti* gene function was collected by searching VectorBase gene IDs on OrthoDB(Kriventseva et al. 2018).

### Statistics

We tested for differences in allele frequency between treatments using generalized linear models that were applied to replicate level major and minor allele counts. We did this in R version 3.2.2 (http://www.r-project.org/). We fitted these single-SNP models using the glm() function in R and assumed a binomial error structure. To aid interpretation, we conducted these analyses in a pairwise fashion, analysing differentiation between all possible pairs of selection treatments (that is, high versus low, high versus random and low versus random populations). To assess the genome-wide significance of these models, and to account for the P value inflation that occurs in single-SNP analyses of evolve and resequence data, we estimated an empirical significance threshold based on exhaustive permutation of our experimental data(Jha et al. 2016). We estimated a permutation-based P value threshold that corresponded to a genome-wide FDR of 5% by re-running our genome scan on all possible permutations of our pairwise contrasts between high and low, high and random and low and random populations. In each case, there were nine possible unique permutations excluding the observed arrangement of the six replicate populations. For each permuted dataset, we refitted our linear model to all SNPs and estimated the number of significant SNPs.

We performed a Singular Enrichment Analysis (SEA) on the AgriGO v2 website using the 61 *A. aegypti* genes associated with *Wolbachia*-mediated dengue blocking listed in the Supplementary File 1. We used the AaegL3.3 reference genome and performed a Fisher test, correcting for multiple comparisons using the Hochberg (FDR) method. We set the significance level to 0.05 and the minimum number of mapping entries to 5. We set the gene ontology type to Generic GO_slim, which includes a reduced version of the terms in the whole GO terms.

### Data availability

Sequence data is available via the European Nucleotide Archive (accession number: PRJEB33044).

---

**Supplementary Table 1.**
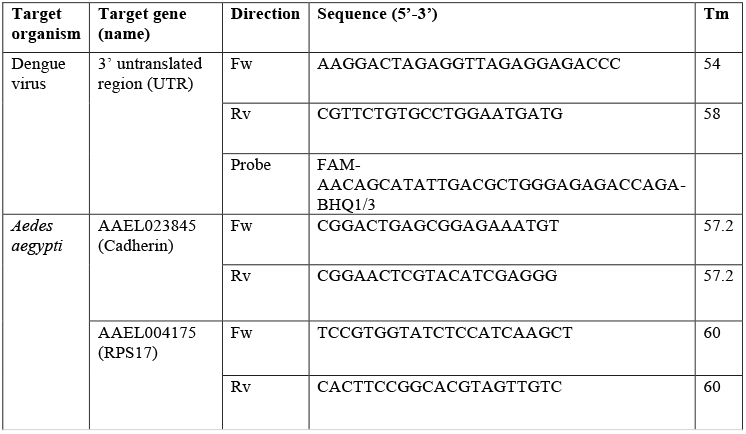
Primers and proves used.

## Notes

https://www.nature.com/articles/s41564-019-0533-3

